# Endocisternal interfaces for minimally invasive neural stimulation and recording of the brain and spinal cord

**DOI:** 10.1101/2023.10.12.562145

**Authors:** Joshua C. Chen, Abdeali Dhuliyawalla, Robert Garcia, Ariadna Robledo, Joshua E. Woods, Fatima Alrashdan, Sean O’Leary, Scott Crosby, Michelle M Felicella, Ajay K. Wakhloo, Patrick Karas, Wayne Goodman, Sameer A. Sheth, Sunil A. Sheth, Jacob T. Robinson, Peter Kan

## Abstract

Minimally invasive neural interfaces can be used to diagnose, manage, and treat many disorders with substantially reduced risks of surgical complications. Endovascular neural interfaces implanted in the veins or arteries is one approach, but it requires prescriptions of anti-thrombotic medication and are likely not explantable after endothelialization. More critically, the approach is limited by the small size and location of blood vessels, such that many important cortical, subcortical, spinal targets cannot be reached. Here, we demonstrate a chronic endocisternal neural interface that approaches brain and spinal cord targets through inner and outer cerebral spinal fluid (CSF) spaces. These spaces surround the nervous system and lack the tortuosity of the circulatory system, giving us access to the entire brain convexity, deep brain structures within the ventricles, and the spinal cord from the spinal subarachnoid space. Combined with miniature magnetoelectric-powered bioelectronics, the entire wireless system is deployable through a percutaneous procedure. The flexible catheter electrodes can be freely navigated throughout the body from the spinal to cranial subarachnoid space, and from the cranial subarachnoid space to the ventricles. We show in a large animal model that we can also reposition the recording and stimulation electrodes or explant the neural interface after chronic implantation. This enables applications in therapies that require transient or permanent brain/machine interface such as stroke rehabilitation and epilepsy monitoring and opens a new class of minimally invasive endocisternal bioelectronics.

## Main

Interest in neurotechnologies that interface with the central nervous system to diagnose and treat diseases is accelerating.^1–4^. As these technologies mature, applications expand beyond simple therapeutics toward more targeted, personalized therapies, and even neural prosthetics enabled by brain machine interfaces (BMIs)^5–7^. A major barrier to entry for these technologies is the ability to interface with our nervous system in a minimally invasive manner which doesn’t damage existing healthy tissue. For example, it is estimated that adoption of deep brain stimulation (DBS) for Parkinson’s Disease (PD) is limited to only about 10% of patients because of the perceived invasiveness and risks associated with procedures such as stereotactic brain surgery required to place electrodes in the brain^8^. Traditional neural implants that interface with the central nervous system (CNS) often rely on penetrating electrodes to access deep brain structures and require invasive surgeries such as craniotomies and burr holes^9–11^.

Recently, innovative surgical approaches via the circulatory system have enabled less invasive implantation of devices that can interface with the nervous system^12^. By inserting catheters through the jugular vein in sheep, somatosensory evoked potentials were measured through the superior sagittal sinus, and more recently, the safety and efficacy of this approach has been demonstrated in humans^12,13^. Additionally, flexible microfabricated electrodes have been developed to navigate sub-millimeter vessels in rodent models for acute neural recordings^14^. However, this method requires a regimen anti-thrombotic medications, and the endothelialization of vascular implants makes it challenging to explant any devices^15^. The tortuous nature of the vasculature and the small caliber of the blood vessels in the brain and spinal cord make it difficult for robust and chronic electrodes to navigate to end targets in the central nervous system, making most cortical, subcortical, deep structures, and spinal cord inaccessible.

We hypothesized that the ventricular system and subarachnoid space could provide an alternative that would allow increased access to the brain convexity and spinal cord and avoid the need for anti-thrombotic medication. Like the circulatory system, the ventricular system and subarachnoid space that holds 150 mL of cerebrospinal fluid (CSF) is tightly coupled with the CNS. The CSF has long been shown to play a crucial role in CNS homeostasis, ensuring mechanical protection, and enabling communication between the CNS and vascular system, lymphatic system, peripheral nervous system, and immune system^16^. CSF is primarily produced in the choroid plexi within the lateral, third and fourth ventricles. It then flows from the ventricular system to the cerebral and spinal surface (subarachnoid space) through the foramen of Luschka and Magendie and is reabsorbed into the dural venous sinus through arachnoid villi. Through this network, this space is in contact with major CNS therapeutic targets and BMI targets such as deep brain structures, brain cortex, and spinal cord. This is also a space familiar to patients and neurosurgeons as many disorders are currently treated using the CSF space and catheter technology (e.g. hydrocephalus, spasticity, and back pain)^17^.

To fully take advantage of these minimally invasive surgical approaches, it is important to reduce the size of the bioelectronic implants. There has been significant progress in miniaturization by developing battery-free devices that rely on wireless power transfer. These techniques that use radio frequency, magnetic fields, and ultrasound enable implants to be miniaturized down to the millimeter scale and operated at centimeter depths within the body^18–27^ These millimeter scaled devices enable the system to also be fully implanted with a minimally invasive percutaneous procedure. Recently, there have been demonstrations of devices that interface with the CNS through CSF such as flexible microelectrodes implanted through surgical holes in the skull into the subarachnoid space^28,29^. Minimally invasive approaches involving implanting electrodes into the epidural space of the spinal column have also been explored^30^. However, the ability to deploy a full system that has access to multiple CNS sites such as brain and spinal cord simultaneously through a minimally invasive procedure and being able to wirelessly stimulate and record these neural structures over extended periods of time has not been reported previously.

Here we present a minimally invasive endocisternal interface (ECI) that accesses multiple targets (cortical surface, deep brain structures, and spinal cord) through the ventricles and subarachnoid space. We also show simultaneous stimulation and recording from both the brain and spinal cord through a minimally invasive percutaneous lumbar puncture. Furthermore, we demonstrate this entire system can be implanted percutaneously by coupling the catheter electrode with a miniaturized implantable pulse generator using magnetoelectric (ME) wireless data and power transfer technology.

## Results

### Endocisternal neural interface enables minimally invasive access to entire central nervous system

We developed a wireless system and a minimally invasive, percutaneous surgical approach that takes advantage of the subarachnoid space to guide catheter electrodes to electrically interface with the central nervous system. In this endocisternal context, the cisternal space is defined as the reservoirs filled with CSF and encompasses both subarachnoid and ventricular spaces of the spinal cord and brain. This approach enables an extremely flexible deployment of the system to a multitude of targets through a simple lumbar puncture (Fig 1A,B). To test this approach, we navigated the spinal and cranial subarachnoid space in human cadaver models. Entering a catheter through the cervical spinal subarachnoid space, we were able to guide a microcatheter (0.6 mm distal tip) into the frontal convexity as well as into the third ventricle in a human cadaver (see methods) under image guidance using cisternography (Fig. 1 B). As can be seen on a cisternogram, the catheter path, highlighted in yellow, passes anterior to deep brain structures like the medulla and pons prior to reaching the 3rd ventricle by traversing its floor (Fig. 1B). Additionally, to guide the development of endocisternal neural interfaces, we characterized the critical dimensions for which an implant could fit within this space. From a lumbar puncture in the lower back, the width of the subarachnoid space falls within the millimeter regime, indicating that mm to sub-mm interfaces could be moved throughout this space^31,32^. Measured across 10 patients, the smallest dimension was found in the cervical spinal region with a measured dimension of 3.93 +/- 1.28 mm. The subarachnoid space remains fairly constant through the lumbar, thoracic, and cervical spinal areas before opening up in the craniocervical junction and narrowing again into the subarachnoid spaces of the brain (Fig 1C, Supplementary Fig. 1). 3D reconstructions of the CSF space showed the mm-sized navigable volumes along with a representative path of the catheter to reach the frontal cortex of the brain (Supplementary Fig. 2, Supplementary Fig. 3). In addition to the human cadaver studies, we validated the ECI in a clinically relevant large animal model. Ovine models have been shown to be an excellent representation for human spinal research^33,34^. Thus, we tested our neural stimulation and recording system in the central nervous system of sheep models interfacing the wireless implantable pulse generator, MagnetoElectric BioImplanT (ME-BIT), with the catheter electrode (Fig. 1D) (n = 12, 10 acute and 2 survival studies).

**Figure 1.**
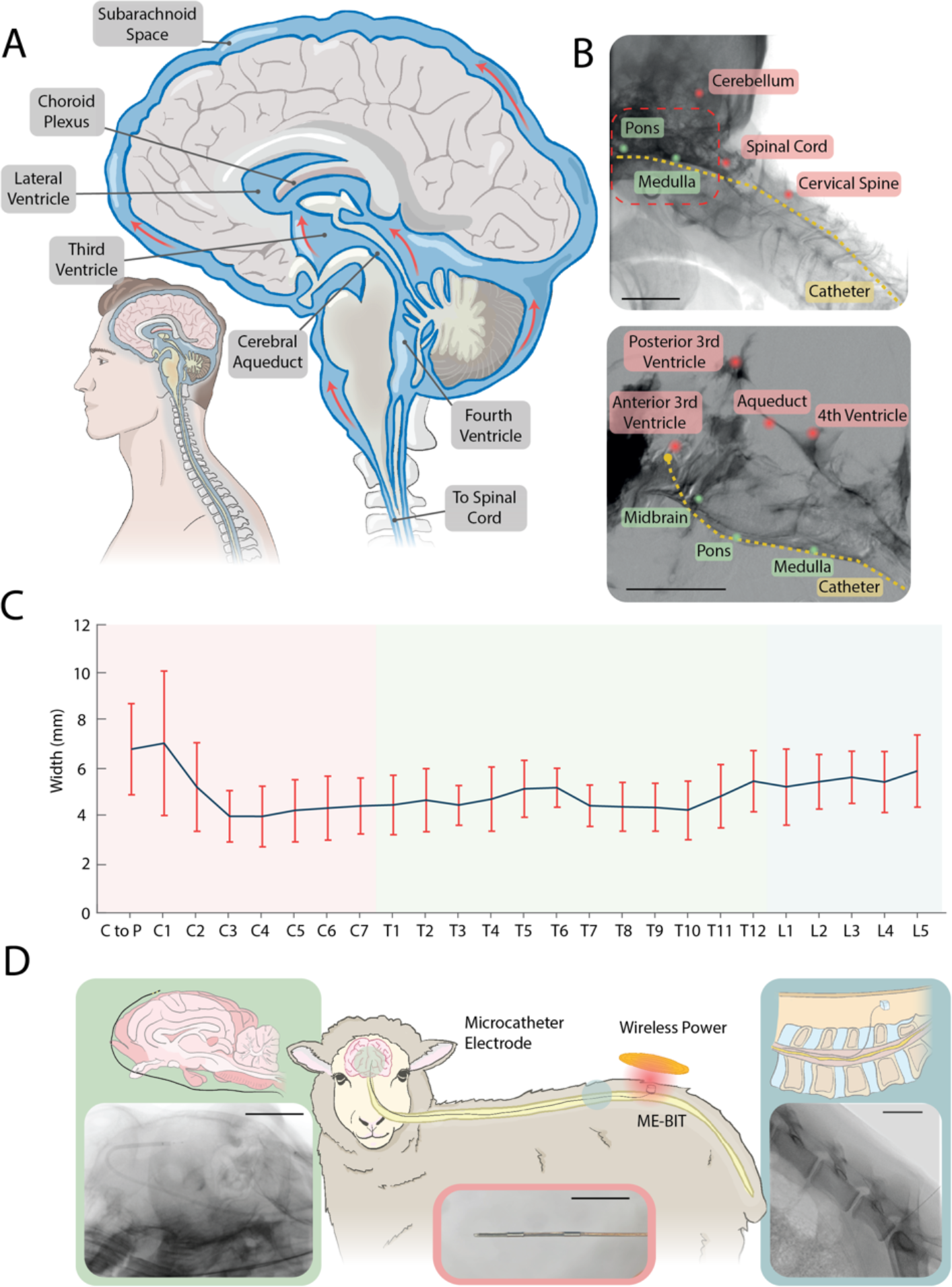
Endocisternal neural interface in human and sheep models: Schematic showing the cisternal space surrounding major neural tissue including the spinal cord and brain. Different navigation pathways can be taken to guide catheter electrodes to reach different therapeutic targets. B. Third ventricular cisternogram in human cadaver models showing the path and guidance of a catheter into the third ventricle along with major anatomical features. Bottom is a zoomed image of the top. (scale bars: top = 50 mm, bottom = 25 mm). C. Width of subarachnoid space in the spinal canal towards deep brain measured across 10 patients with MRI. Measurements are an average of the anterior and posterior subarachnoid spaces. D. Schematic of ECI deployed in a sheep model with implanted magnetoelectric pulse generator (ME-BIT) and wireless power transmitter. Left schematic shows a catheter navigated to the frontal cortex of the sheep brain and corresponding X-ray below with the catheter visible (scale = 50 mm). Center image shows a magnified image of the two-channel catheter electrode. Right schematic shows the pulse generator implanted in the lumbar region of the sheep with the catheter electrode entering subarachnoid space along with a corresponding X-ray of a lumbar puncture with a catheter (scale = 50 mm).

### Wireless stimulator demonstrates coordinated and multi-site stimulation and recording in large animal model

Using a millimeter-sized battery-free implantable stimulator we demonstrate multi-site electrical stimulation and recording through the ECI. We take advantage of recent developments in magnetoelectric materials, which are thin films that convert magnetic fields to electric fields, to engineer miniaturized battery-free bioelectronics to serve as the implantable pulse generator^35^. By using a magnetic field, we can not only send power and data into the implant to have programmable stimulus pulses with varying amplitudes up to 14.5 V, pulse widths, and duty cycles, but we are also able to take advantage of recently developed magnetoelectric backscatter to also wirelessly record physiologically relevant signals through this endocisternal neural interface^36^ (Fig. 2D, Supplementary Fig. 4). Briefly, the implant consists of a printed circuit board coupled with a magnetoelectric films and a bias magnet to optimize the power transfer on the films. These components are assembled inside a glass capsule and sealed with medical grade epoxy with stimulation and recording leads coming out of the package (Fig. 2D). The ME-BIT is placed in a small pocket in the back of the animal and electrically wired to the catheter electrode that is inserted into the subarachnoid space. Additionally, the system includes the wireless transmitter that uses a litz wire pancake coil coupled with recording coils on a PCB (see methods Fig. 2D).

**Figure 2.**
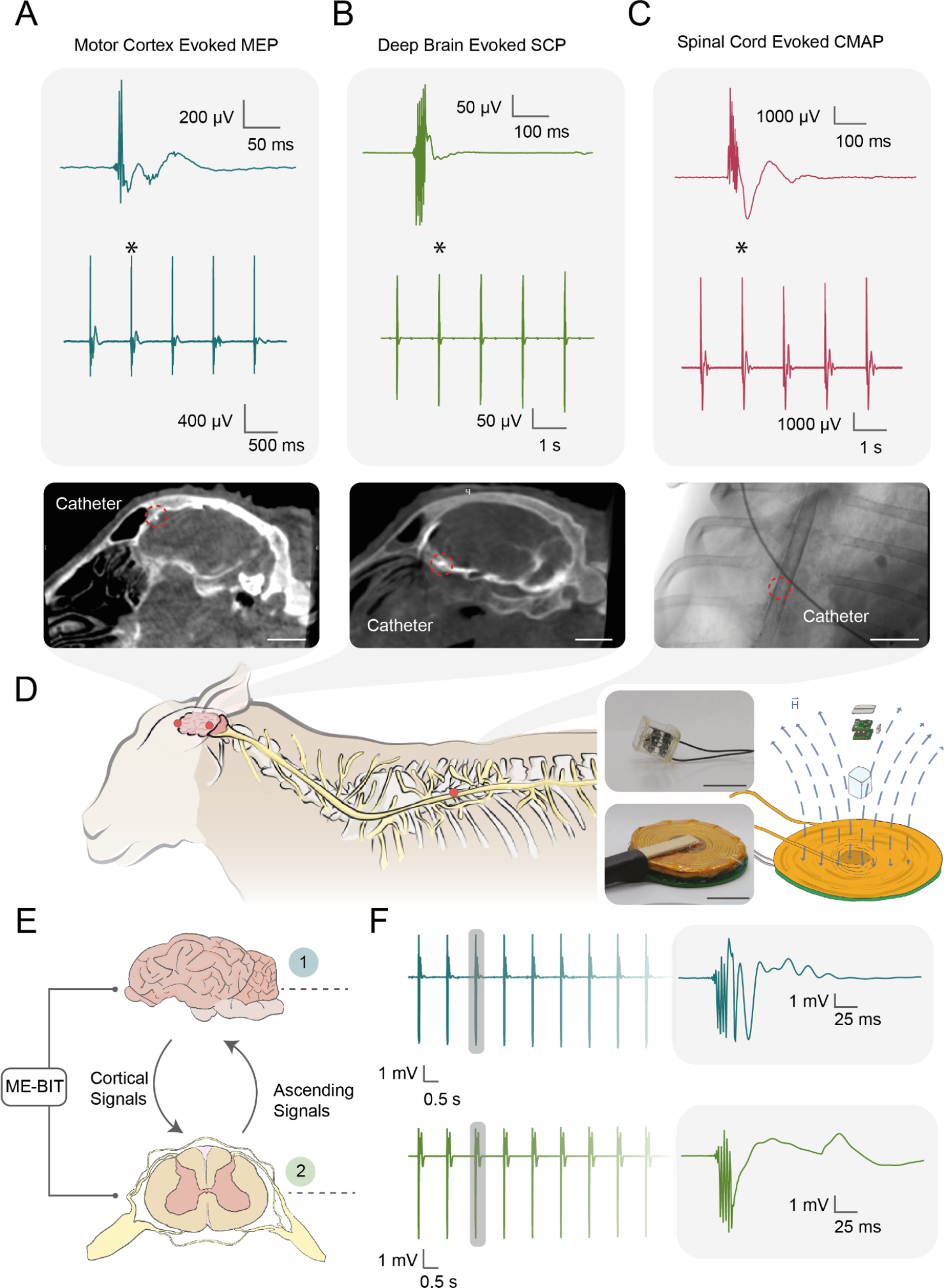
Wireless neural stimulation through endocisternal interface in sheep: **A** Recordings of motor evoked potentials recorded from the hind leg of the sheep from stimulation in the motor cortex. Corresponding CT shows the catheter tip circled in red at the frontal cortex of the sheep brain. (scale = 20mm). **B**. Electrophysiological recordings of spinal cord potentials that were evoked through stimulation of deep brain structure (thalamus). Associated CT shows catheter position where the tip is circled in red. (scale = 20 mm) **C**. Stimulation of the spinal cord results in recorded compound muscle action potentials in surrounding back muscles. X-ray shows catheter position inside the subarachnoid space around the spinal cord. (scale = 50 mm) D. Schematic of the sheep and positions of catheter electrode and neural stimulation. Images on the right shows the magnetic field transmitter and magnetoelectric powered implant and schematic of the wireless system. (scale - implant = 1 cm, scale – transmitter = 3 cm) **E**. Schematic showing the synchronized stimulation of brain and spinal cord with the magnetoelectric powered implant where cortical and ascending neural signals are sent between the two structures. **F**. Time locked recordings of simultaneous brain and spinal cord stimulations with surface needle electrodes showing resulting CMAPs.

Due to the flexibility of deployment, we navigated the catheter electrodes starting at the lower back up the spinal canal and into the brain. We were able to stimulate throughout the spinal cord from lumbar up to cervical regions. Powering the electrode with the ME-BIT and applying a 14.5V, 250 us pulse width monophasic pulse train of 10 pulses at 0.8 Hz activated the spinal cord. We observed muscle contractions and measured compound muscle action potentials (CMAPs, Fig 2C, Supplementary Fig. 5A) with EMG needles on the back of the sheep. The catheter was further navigated through the craniocervical junction to enter the brain.

Without any burr holes or craniotomies, we were able to apply the same electrical stimulation waveform of 14.5V at 2 Hz to stimulate the motor cortex where we observed muscle contractions in the hind leg of the sheep and measured corresponding MEPs (Fig. 2A, Supplementary Fig. 5B). Additionally, stimulated deep brain structures (thalamus through ventricular surface) at 0.8 Hz and record downstream activation of the central nervous system with spinal cord potentials. In fact, by increasing the number of stimulation pulses, we confirmed that we indeed elicited a central response through observations in latency shifts as we increased stimulation amplitude^37^ (Supplementary Fig. 6). Furthermore, we simultaneously fit two catheter electrodes with diameters 0.6 mm within the subarachnoid space with the first electrode navigated to the cortex of the brain and the second catheter electrode on the spinal cord in the thoracic region (Fig. 2B). We were able to apply simultaneous electrical stimulation using one ME implantable pulse generator to synchronously stimulate both brain and spinal cord demonstrating simultaneously access. Simultaneous multisite access remains highly relevant for developing neural interfaces that can perform closed loop and coordinated stimulation and recording across different neural structures^38,39^. Recording the local field potentials at the site of stimulation in both the head and leg yielded measurable MEPs and CMAPs where the stimulus artifacts are both time locked (Fig. 3E). Future development of the catheter electrodes would enable multichannel functionality with exposed electrodes along the length of the catheter to reduce the number of catheters implanted in the body.

**Figure 3.**
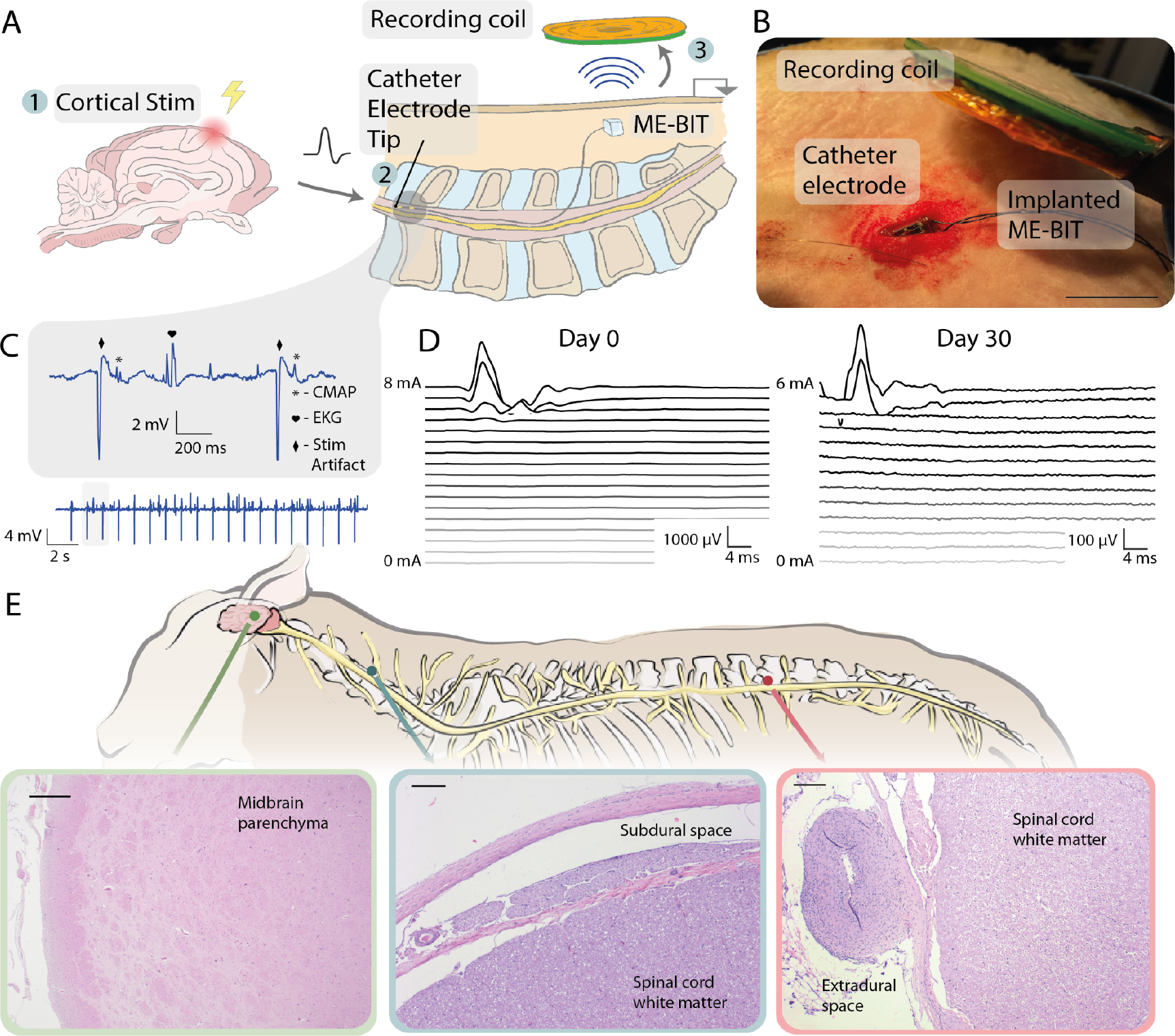
Wireless recording and chronic implantation of endocisternal interface: **A** Schematic of experimental setup to validate wireless recording with the ECI. Frontal cortex of the brain is stimulated with a wireless ME implant through a catheter electrode (1). A central signal is propagated to the spinal cord where it is recorded at the catheter electrode tip (2). The signal is sent wirelessly through magnetoelectric backscatter from the implant to the recording coil (3). **B**. Experimental setup with exposed magnetoelectric implant with stimulation leads (black wires). The catheter electrode implanted through a lumbar puncture (silver wire) and the wireless transmitter is shown. (scale = 1.5 cm) **C**. Wirelessly recorded electrophysiologic signal through ME backscatter from the catheter electrode in the subarachnoid space. **D**. Muscle activity recordings with increasing amplitude comparing stimulation threshold at Day 0 and at Day 30 in a sheep model. **E**. Histology from a sheep implanted with the ECI. The animal was implanted and stimulated at day 0, stimulated again at day 30 and the device explanted, and the sheep was sacrificed one week later. (scale left = 500 um, middle = 200 um, right = 200 um).

We further tested the functionality of this endocisternal neural interface to wirelessly record electrophysiologic signals. We implemented a wireless backscatter protocol with 3 bits for downlink and 8 bits for uplink where the magnetic field is pulsed, and the magnetoelectric ringdown characteristics are measured while the field is off. The data ‘0’ and data ‘1’ are encoded by changing the applied electric load to the magnetoelectric films (Supplementary Fig. 4B, see methods)^35^. We implanted two catheter electrodes in the body, one in the brain over the motor cortex connected to a ME stimulator, and the second in the spinal cord connected to a separate ME recording implant. We then electrically stimulated the motor cortex with 1 Hz pulses through the initial catheter and hypothesized that we could record the downstream electrophysiologic signals with the catheter placed in the spinal cord (Fig 3A/B). Indeed, we observed CMAPs associated with muscle contractions and were able to see extraneous signals such as heart activity and the stimulus artifact (Fig. 3C). Due to current limitations in the sampling frequency of magnetoelectric backscatter and gain of the wireless recording circuit, we also used a benchtop recording system to record smaller magnitude signals such as spinal cord nerve potentials where we observed D-wave activity upon brain stimulation (Supplementary Fig. 7). Additionally, using the benchtop system, we could record EEG from the frontal convexity of the brain. (Supplementary Fig. 8A/B).

### Chronic implantation shows long term viability and explantability

When implanting for more than 30 days in the sheep model we found stable stimulation thresholds on day 0 and day 30 suggesting good chronic performance of the endocisternal neural interface (n=2). With the catheter electrode placed in the frontal convexity, by increasing the amplitude of a single stimulation pulse with a 250 us pulse width, we begin to observe muscle contraction at around a stimulus current of 6.5 mA (Fig 3D, see methods). At day 30, we see a similar stimulation threshold of around 5.5 mA which indicates no noticeable changes in the electrical connectivity of the interface to the tissue. The catheter also remained stable in position. Differences in the amplitude of the recorded muscle activity are likely due to variation in placement of EMG recording needles. In addition to surviving the animal for 30 days with no neurological deficits observed, we also demonstrated explantabilty of the ECI and survived the sheep for an additional week, again with no neurological deficits. This unique property of the endocisternal interface will allow this to be used in transient bioelectronic therapies where explantability is required^40^. A week after explantation, the sheep was sacrificed and upon conducting histology, we found no subdural, subarachnoid, or parenchymal inflammation or damage at any level in the spinal cord, midbrain, cerebrum, or cerebellum (Fig. 3E). There was only some mild extradural inflammation found in the lumbar region which could be a result of the lumbar puncture (Supplementary Fig. 9). Otherwise, spinal cord, brainstem, cerebrum, and cerebellum were all normal after the 30-day chronic implantation and explantation. In another sheep that was survived for a month, we also found no inflammation in the brain or spinal cord with only minor cervical inflammation in the subdural space and minimal inflammation in the leptomeninges of the midbrain which is within expectations for any implant. (Supplementary Fig. 10).

## Discussion

The preliminary feasibility and safety of endocisternal navigation and minimally invasive stimulation and recording from ECI implants shown here in acute and survival animal studies suggests the potential of future minimally invasive ECI implants and transient therapies where the ECI implant can be safely removed after several weeks^40^.

While we developed an initial catheter prototype, further exploration of the catheter interface and other geometries such as mesh electrode arrays would enable multiplexed stimulation through the subarachnoid and ventricular spaces. Our initial prototype has two electrode channels which enable bipolar stimulation in one channel or monopolar stimulation with two channels and a reference electrode on the implantable pulse generator, but multi-channel arrays are common in strip electrodes like those used in spinal cord stimulators^41^. While we were able to explant our device after more than one month in a large animal, future work can explore longer chronic implantation and explantation of different electrode interfaces. A key advantage of these endocisternal neural interfaces is the ability to access both spinal cord and brain structures simultaneously, which is also important for other applications such as paralysis rehabilitative therapies where coordinated stimulation and recording between brain and spinal cord can induce plasticity and recovery^38,39^.

The ease for which these endocisternal neural interfaces can reach the entire brain convexity along with deep brain structures through the ventricles enables neuromodulation of difficult to reach areas in through a simple lumbar puncture. Other known access to the subarachnoid space such as C1/2 and suboccipital punctures can be explored in future experiments. Suboccipital puncture can potentially access the dentate nucleus to enhance stroke rehabilitation through deep brain stimulation^42^. As demonstrated in this manuscript, we were able to not only access the motor cortex which has implications for brain machine interfaces to restore motor functions, but also deep brain structures like the thalamus through the 3^rd^ ventricle which can be used to treat movement disorders^43^. Additionally, this technology allows for easier access to certain brain areas such as the orbitofrontal region (potential target for refractory psychiatric disorders) where traditional surgical access is difficult^44,45^.

Future studies would be required to develop navigation under multi-modality image guidance (e.g. the merge of MRI and myelogram) and further testing can be done to provide more targeted and focal stimulation as this technology matures. An access system specifically designed for endocisternal navigation is also needed to optimize access and navigation in the endocisternal space. The minimally invasive surgical approach shown here may help increase the adoption rates for these implants as they become more minimally invasive. Especially as more neurotechnology is primed for translation, it becomes more imperative to minimize the risk and recovery for patients to increase adoption. With interfaces like these, that can access brain and spinal cord with a catheter, we can envision neural interfaces becoming as ubiquitous as other bioelectronic implants such as cardiac pacemakers or cochlear implants.

## Methods

### Human cadaver navigation

Under direct visualization, a 4F Terumo slender sheath was directly inserted into the dorsal cervical spinal subarachnoid space of the cadaver. 20 cc of iodinated contrast was infused through the sheath to provide a cisternogram and myelogram for navigation. Through the sheath, a Synchro 0.014” microwire (Stryker, Kalamazoo, Michigan) is used to navigate a SL10 microcatheter (Stryker, Kalamazoo, Michigan) from the cervical subarachnoid space to the intracranial subarachnoid space by staying ventral to the spinal cord and midline under fluoroscopic guidance (Artis Zeego, Siemens). Once intracranially and as described previously, the catheter was navigated into the third ventricle through traversing the floor of the third ventricle behind the clivus (Fig. 2c, d)^46^. Third and fourth ventriculograms were performed through the microcatheter.

### Magnetoelectric implantable pulse generator

The implantable pulse generator used in this study is described in detail in Millimeter-sized battery-free epidural cortical stimulators^35^. In brief, the implantable pulse generator is composed of a custom printed circuit board (PCB), two 7.5x3mm magnetoelectric films (resonant frequency 218kHz), and a glass enclosure. The leads are connected to the glass device using a conductive epoxy attachment to sputtered electrodes. The device is programmed to output a train of 10 voltage controlled, programmable amplitude, biphasic pulses, at a rate of 500 Hz when triggered by the external transmitter. The external transmitter communicates with the device using pulse-width modulated downlink communication.

### Magnetic Field Transmitter

A driver system was built high-electron-mobility gallium-nitride transistors (GS61008T,GaN Systems) as the H-bridge stage with an optimized magnetic board layout for high frequency and power switching operations^47^. This magnetic field driver was controlled using a custom script through Waveforms where control signals were sent through an Analog Discovery Pro (ADP3450/ADP3250). Together this system drives current through a resonant coil that is wrapped using 18 AWG litz wire (MWS Wire) and resonated with high voltage rated capacitors (WIMA).

### Histology of Explantation of Endocisternal Device

The sheep were stimulated with the ECI on day 0 and survived for 30 days. After the month, the interface was explanted and the sheep are survived for another week before being sacrificed. Tissue slices were taken of both the spinal cord and the brain. Standard Hematoxylin and Eosin (H&E) staining was used to visualize the tissue anatomy.

### Human subarachnoid space measurements

Results of sagittal T2 weighted fast relaxation fast spin echo sequence (FRFSE) MRI images of the cervical, thoracic, and lumbar spinal canal in 10 live human patients, conducted without the administration of intravenous (IV) contrast (1.5 T GE and Siemens and 3T Siemens). Measurement of the spinal canal was performed using Picture Archiving and Communication System (PACS) built in measurement tools. Dimension of the entire spinal canal was measured along with both the posterior and anterior sides of the canal subtracting the spinal cord. The final average between posterior and anterior measurements are reported in the main figure (Supplementary Fig. 11).

### In vivo ovine surgical access for wireless recording and stimulation

The animal procedures were conducted in accordance with regulations of the IACUC (Protocol no. 2110061). Twelves female Western Range sheep, weighing approximately 40-50 kg, received a 7-day acclimation period prior to any procedure. General anesthesia was induced by ketamine (5 mg kg^-1^) and xylazine (0.2 mg kg^-1^) intramuscularly, followed by intubation by veterinary personnel. Mechanical ventilation was maintained under a mixture of oxygen and isoflurane (1-3%). Routine physiological monitoring was performed. Animal was placed in prone position to allow palpation of anatomical landmarks and initiation of the lumbar puncture.

Under fluoroscopic guidance (CIOS Spin 3D, Siemens Healthineers), we access the lumbar subarachnoid space of the sheep with an 18 G spinal needle. Once access is confirmed with spontaneous flow of cerebrospinal fluid (CSF), an 0.018” wire is threaded under fluoroscopic guidance up to the thoracic subarachnoid space. Using the Seldinger technique, the spinal needle is then exchanged for a 5F Glidesheath Slender introducer sheath (Terumo, Somerset, New Jersey). A myelogram is then performed through the sheath to ensure proper placement (e.g. not in the epidural space). A Synchro 0.014” microwire (Stryker, Kalamazoo, Michigan) is used to navigate an Echleon 10 microcatheter (Medtronic, Minneapolis, Minnesota) from the lumbar subarachnoid space to the intracranial subarachnoid space by staying ventral to the spinal cord and midline under fluoroscopic guidance. Once intracranially, the catheter can be navigated along the clivus to the orbitofrontal cortex and around the frontal pole to the frontal convexity (target of stimulation Fig. 1B). Alternatively, it can access the third ventricle through the floor of the third ventricle. Through the third ventricle, the thalamus can be accessed (Fig. 1B). Electrophysiologic recordings were done on a Cadwell IOMax using subdermal needles for the EMG recordings, otherwise the benchtop recording system was connected to the endocisternal catheter electrodes. Additionally, the benchtop system was used for thresholding experiments. Wireless recording was done using the magnetoelectric implant. Two sheep were used for survival chronic implantation experiments and the other 10 sheep were used for acute stimulation and recording experiments. Successful stimulation and recording were done in all 12 of the sheep.

## Supplemental

**Supplementary Figure 1.**
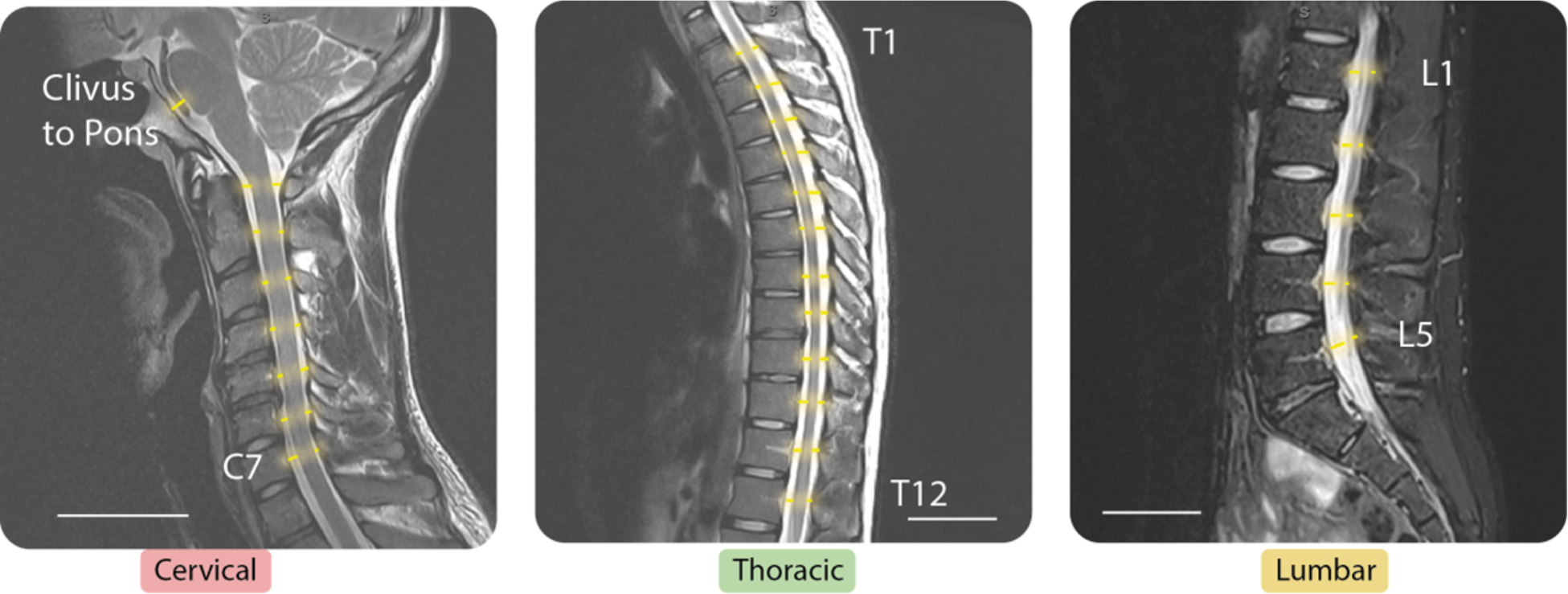
MRI images of human cadavers in cervical, thoracic, and lumbar regions to measure dimensions of CSF space.

**Supplementary Figure 2.**
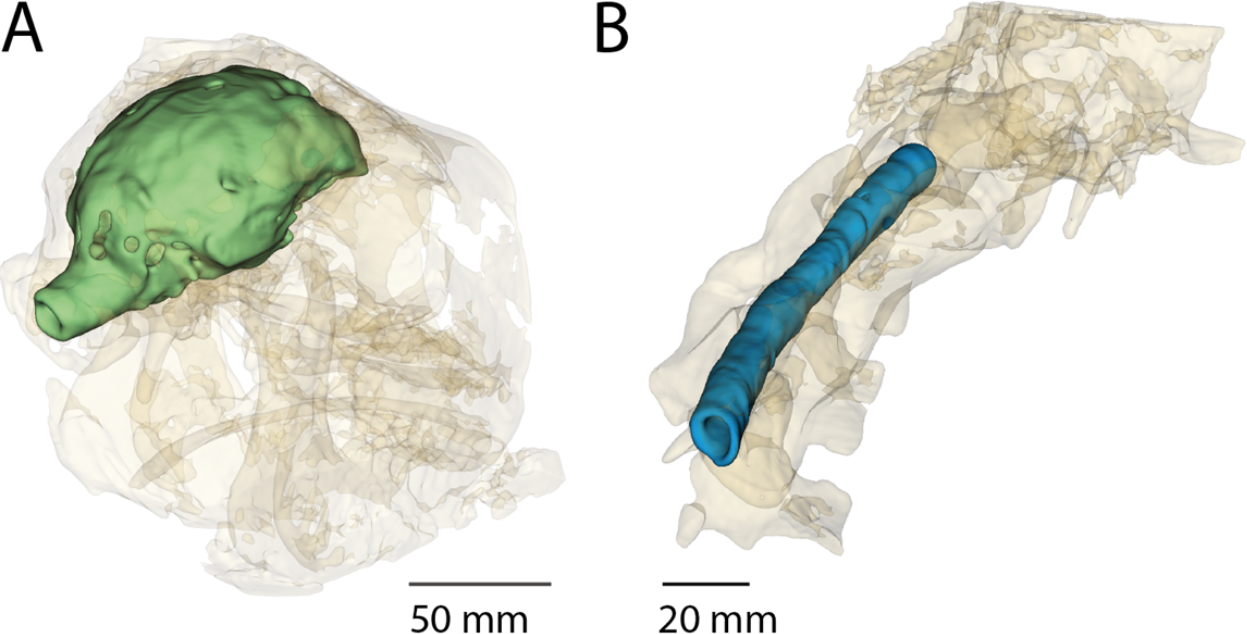
3D reconstruction of the sheep head highlighting the CSF space (green) visualized by injecting contrast through a navigated catheter. B. 3D reconstruction of the spine highlighting the subarachnoid space (blue) surrounding the spinal cord.

**Supplementary Figure 3.**
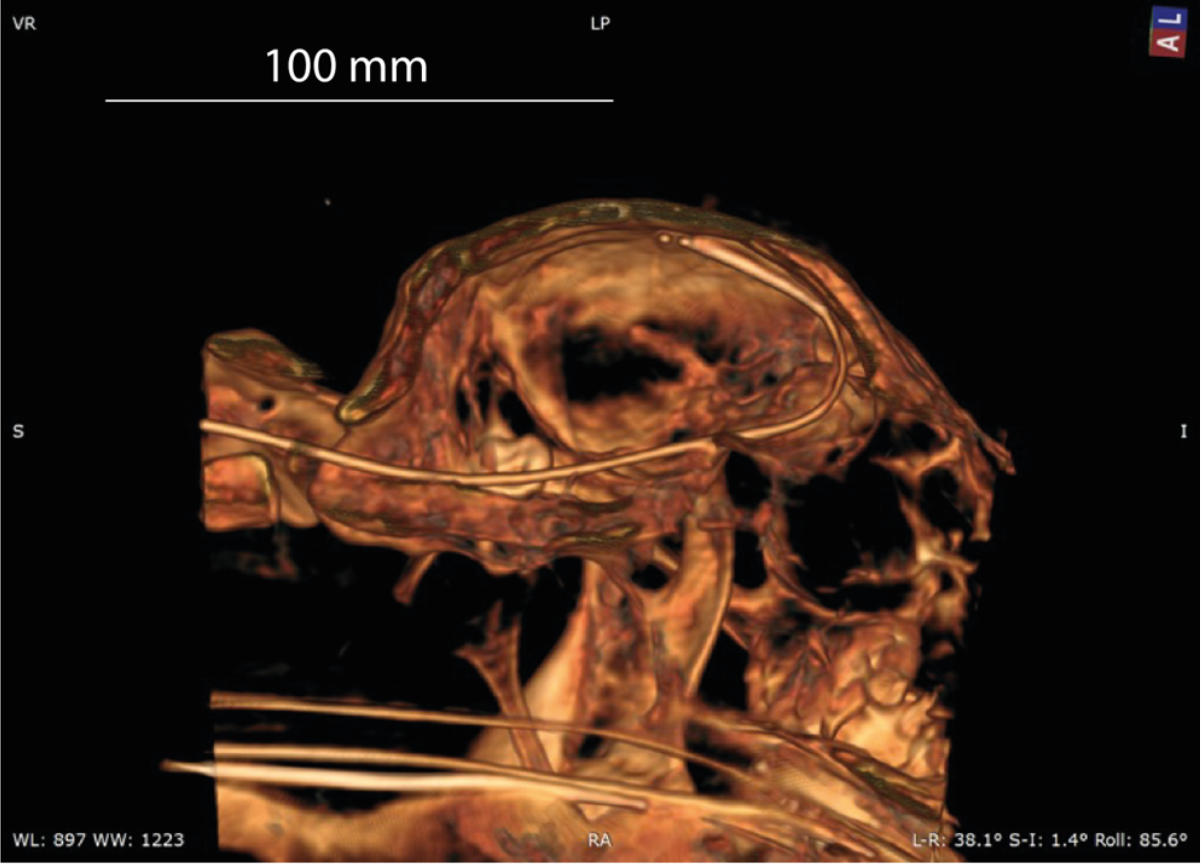
3D reconstruction of the catheter electrode going through craniocervical junction and navigating to frontal cortex of the brain.

**Supplementary Figure 4A.**
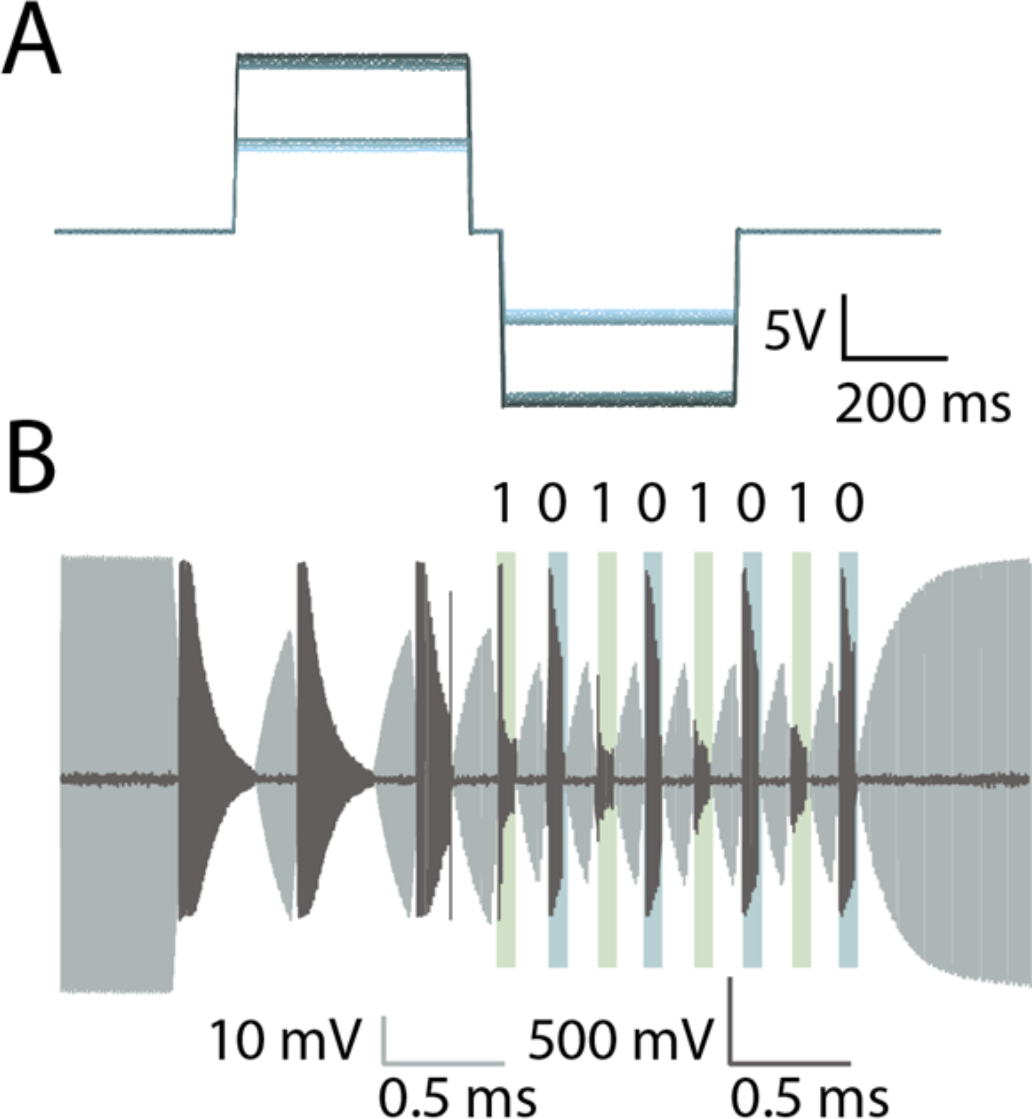
Programmable voltage-controlled stimulation pulses generated by the magnetoelectric wireless implant. B. Sample data uplink where light grey is the transmitter coil current, and dark grey represents the backscattered magnetic field.

**Supplementary Figure 5A.**
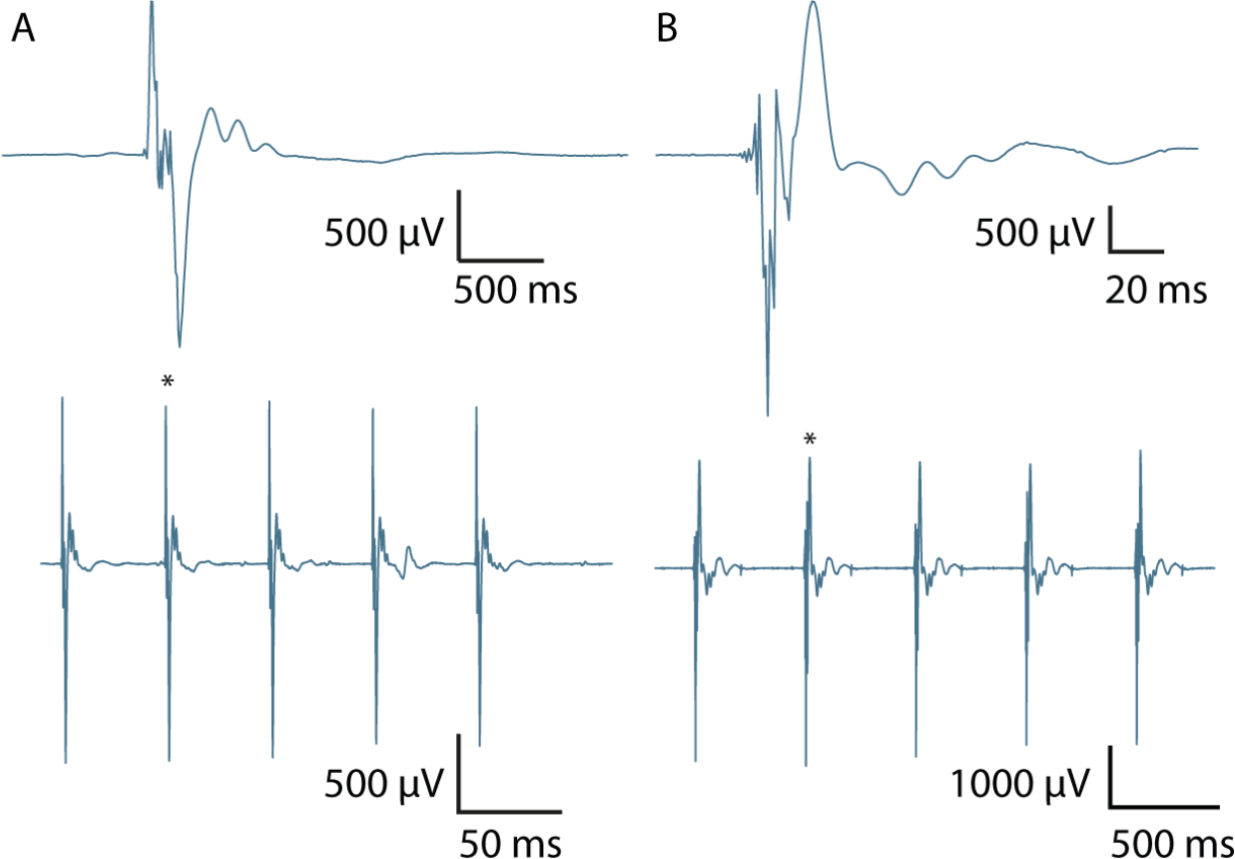
Stimulation with bipolar electrodes in the thoracic region of the spinal cord and recorded CMAPs. **B**. Stimulation with bipolar electrodes in the frontal convexity of the brain and recorded CMAPs on the hind legs of the sheep.

**Supplementary Figure 6.**
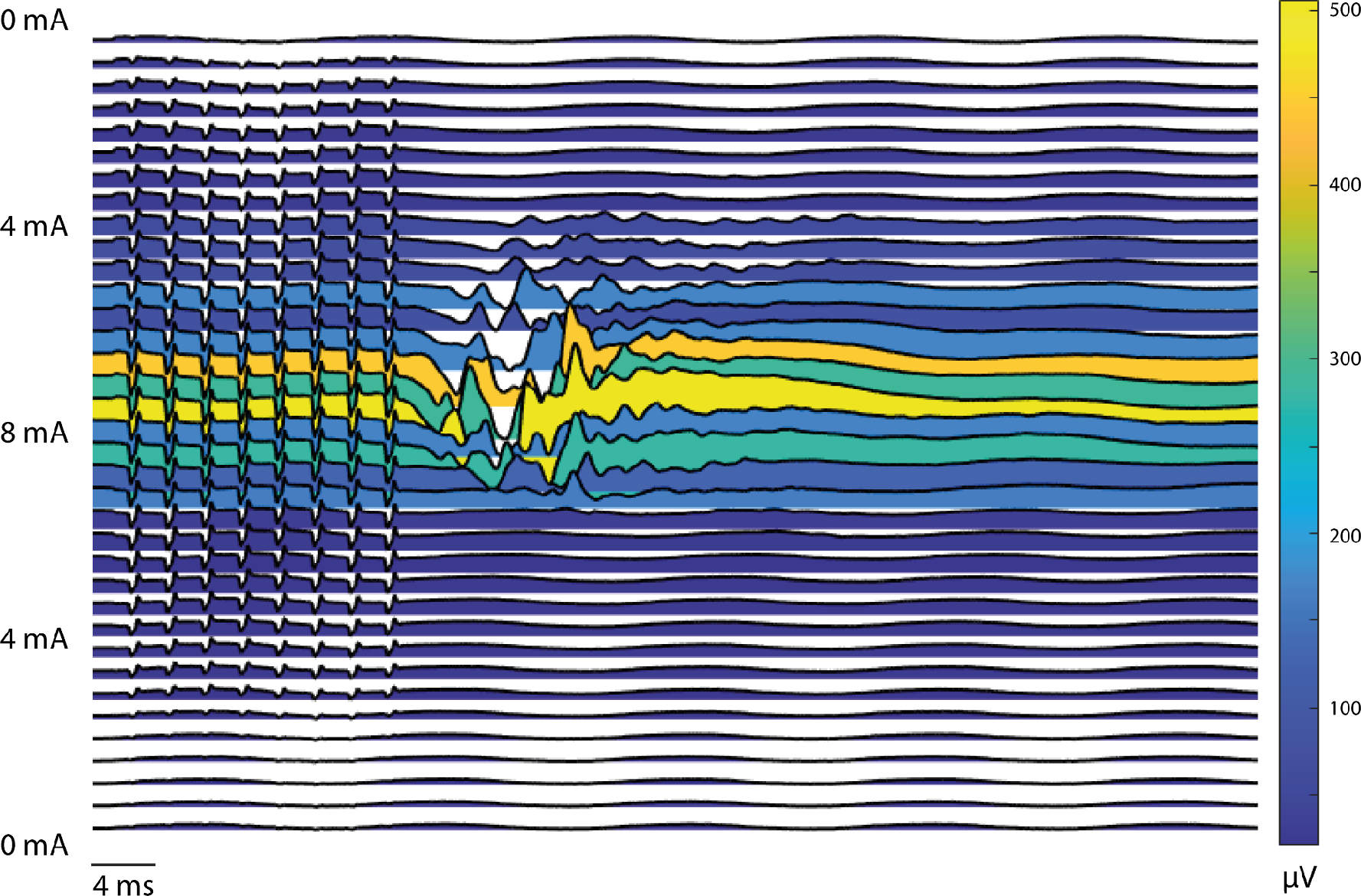
Nine pulse waterfall plot showing central activation with shifting latencies with first increasing stimulation amplitude from 0 mA to 8 mA and then decreasing back down to 0 mA. Color represents the max voltage reached on the trace.

**Supplementary Figure 7.**
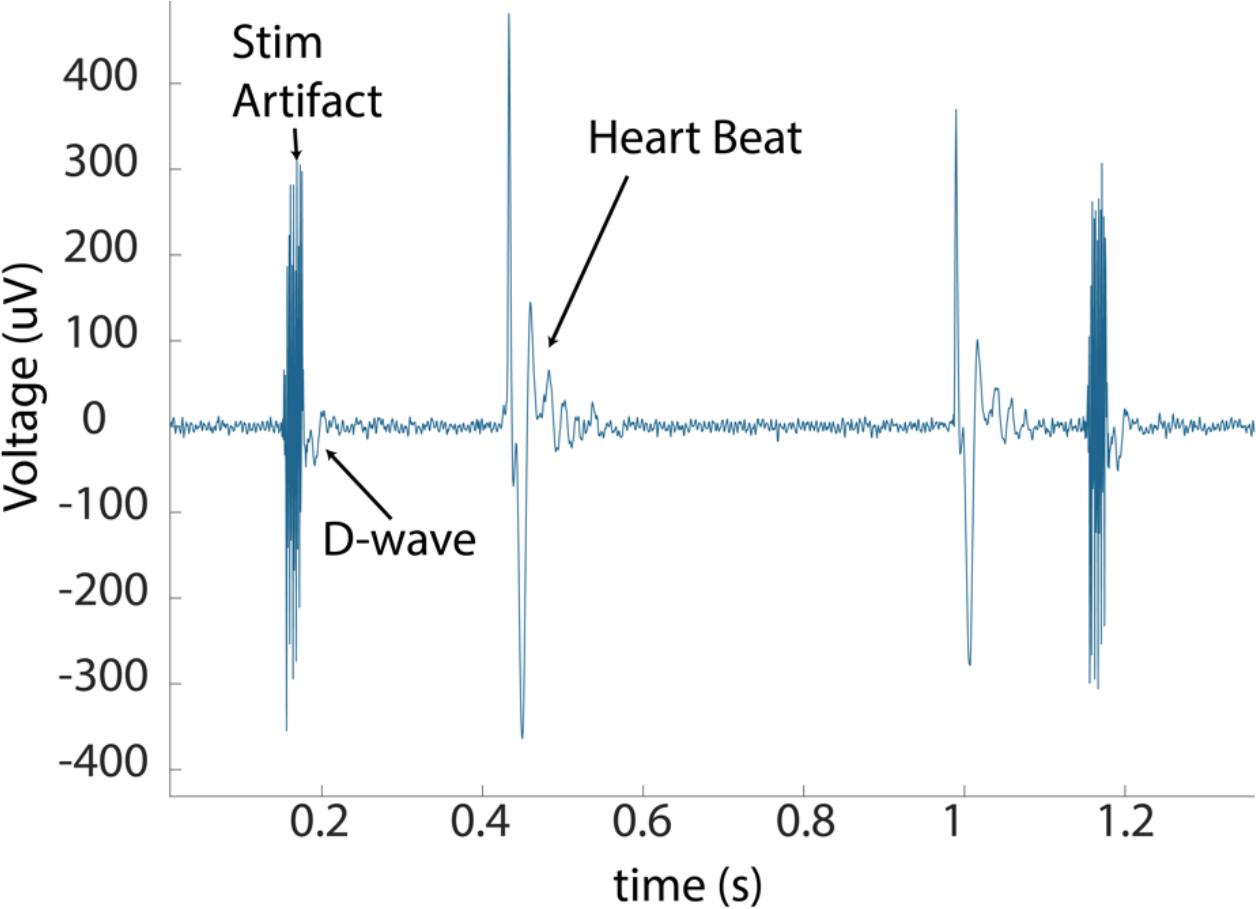
Catheter recording from within the subarachnoid space surround the spinal cord where a D-wave is visible following the high frequency stimulation artifact. EKG is also visible.

**Supplementary Figure 8A.**
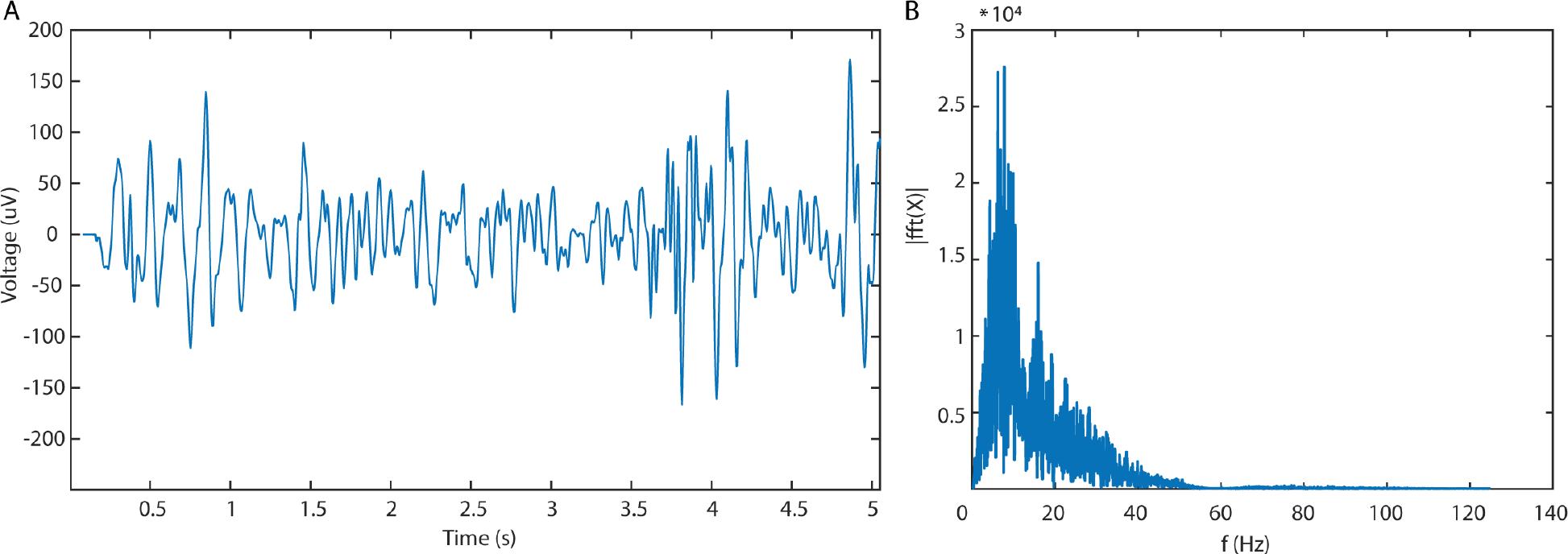
EEG recording through catheter electrode from the frontal convexity of the brain. **B**. Fast fourier transform of the EEG signal to highlight the frequency components of the signal.

**Supplementary Figure 9.**
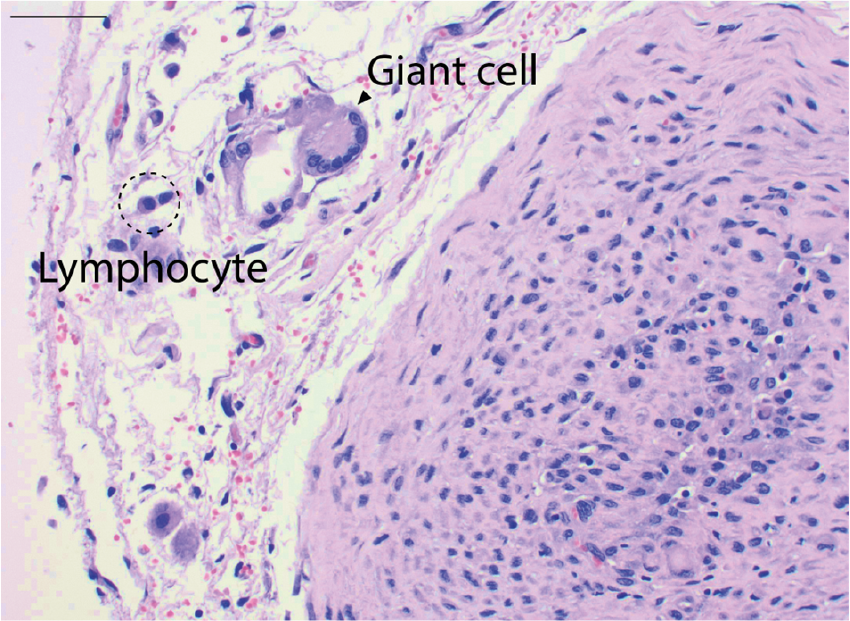
High power image of lumbar epidural reactive vasculature with mild lymphocytic and histiocytic inflammation with rare foreign body-type giant cells.

**Supplementary Figure 10.**
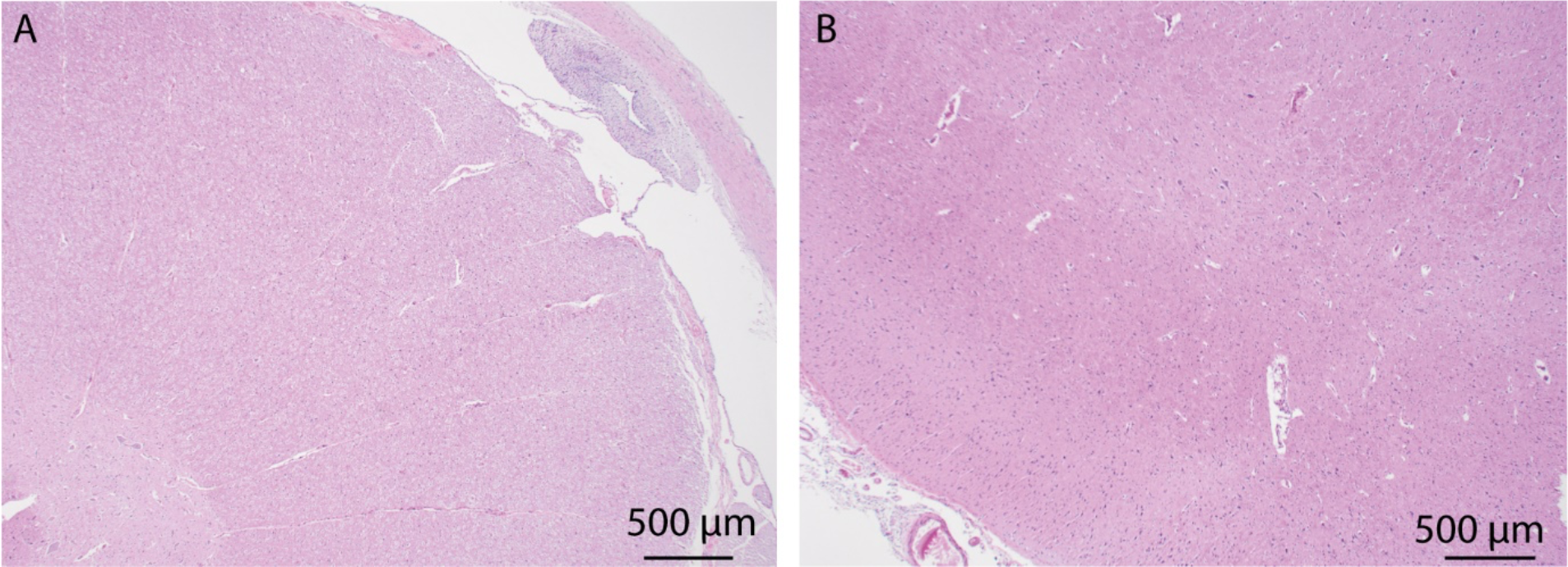
**A** Cervical spinal cord histology in survival sheep 2 showing mild inflammation in subdural space. **B**. Histology of midbrain in survival sheep 2 showing normal parenchyma and only minimal inflammation in leptomeninges.

**Supplementary Figure 11.**
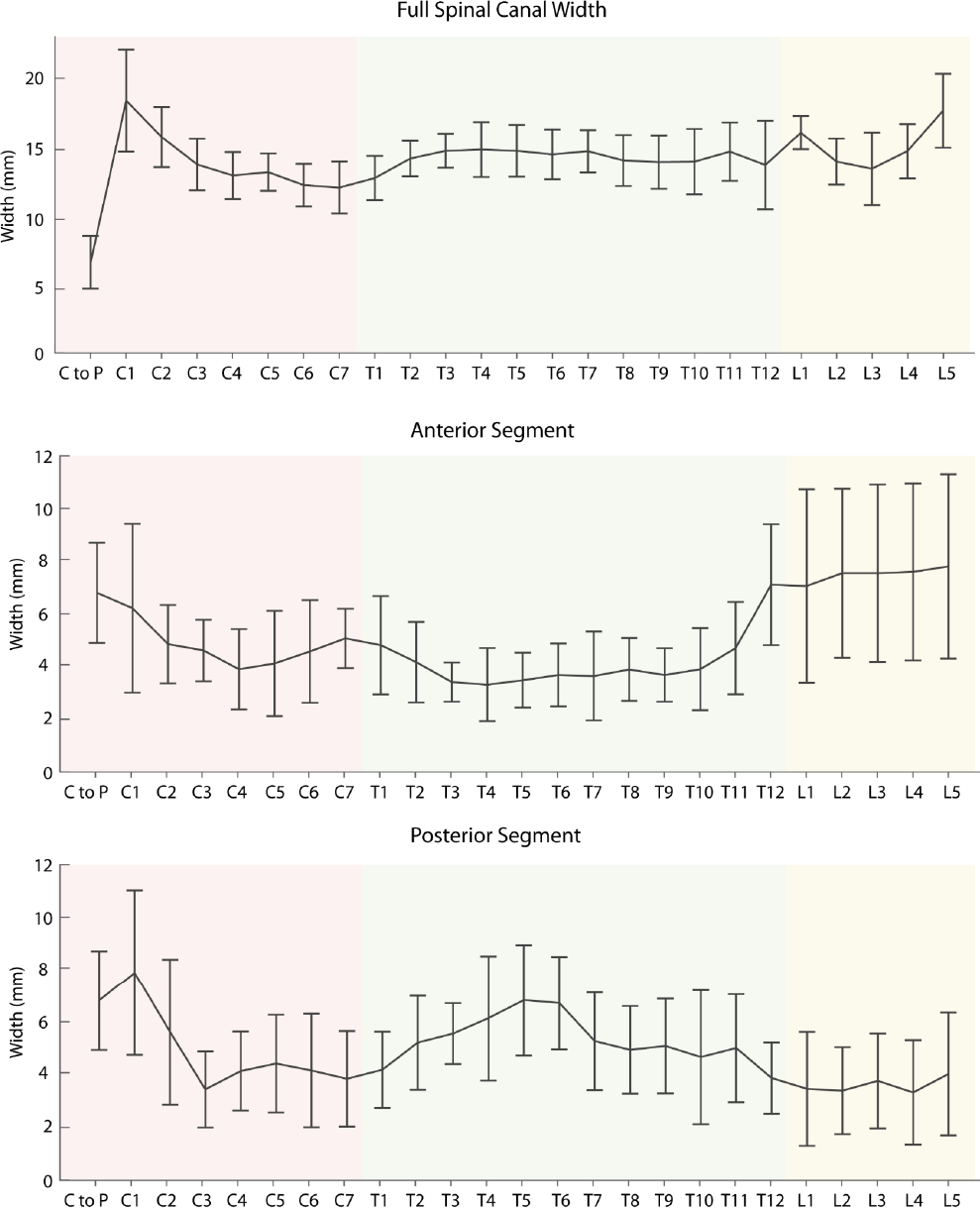
Measurements of spinal canal width taken from the MRIs of human cadavers. Data presented in Figure 1 is the average of the anterior and posterior segments.

## Acknowledgements

We would like to thank Deinde Medical for introducing us to CSF microcatheter navigation and endocisternal techniques for 3^rd^ ventriculostomy in the ovine model.

